# Molecular, physiological and functional features underlying cortical thinning related to antipsychotic medication use

**DOI:** 10.1101/2024.01.05.573095

**Authors:** Lauri Tuominen, Reetta-Liina Armio, Justine Y. Hansen, Maija Walta, Nikolaos Koutsouleris, Heikki Laurikainen, Raimo K.R. Salokangas, Bratislav Misic, Jarmo Hietala

## Abstract

Use of antipsychotic medication is related to thinning of the cerebral cortex, but the underlying mechanisms of this effect remain largely unknown. Here, we investigated potential mechanisms across multiple levels of description, from molecular and physiological factors to whole-brain functional patterns. We first analyzed a single site discovery sample of patients (N=131) with early psychosis for whom antipsychotic related cortical thinning was estimated based on lifetime exposure to antipsychotics. Findings were replicated using data from a large (N≥2168) ENIGMA meta-analysis. We discovered that antipsychotic related cortical thinning is associated with a number of neurotransmitter systems, most notably the serotonin system, physiological measures, and functional networks and neural oscillatory power distributions typical for regions subserving higher cognition. At the functional level, antipsychotic related cortical thinning affects regions involved in executive function and motivation, but not perception. These results show how molecular, physiological, and large-scale functional patterns underlie antipsychotic related cortical thinning.

## Introduction

Antipsychotic medication is the first-line treatment of psychotic symptoms in schizophrenia and other psychotic disorders (1). These medications effectively alleviate positive psychotic symptoms such as hallucinations and delusions (2,3). Long-term use of antipsychotics has been shown to reduce relapses and rehospitalizations, aggressive and suicidal behavior, and overall mortality in individuals with schizophrenia (4–6). In addition to the established clinical side effects such as weight gain and extrapyramidal motor side effects, antipsychotics have a well-documented effect on brain morphology. A number of longitudinal studies including studies in previously drug naïve participants have demonstrated an association between cumulative exposure to antipsychotics and decreased cortical gray matter (7–9). These findings are mirrored in a large cross-sectional ENIGMA meta-analysis that found a similar correlation between reduced cortical thickness and current daily dose of antipsychotics (10). Causal evidence that antipsychotic treatment leads to decrease in cortical thickness has been provided by animal models and at least one recent randomized controlled trial in humans (11,12).

The exact mechanisms by which antipsychotic use leads to decreased cortical gray matter still remain mostly unknown. Citing both preclinical and clinical research, Turkheimer and colleagues recently proposed that antipsychotics might push the cortex into an unsustainable metabolic envelope and increased state of oxidative stress, which would in turn lead to adaptive changes in cortical morphology (13). According to this hypothesis, antipsychotics normalize metabolic changes related to psychosis, however this normalization is ultimately unsustainable and thus leads to cortical remodeling. Importantly, these effects are suggested to depend on efficient mitochondrial function, adequate regional blood supply, functioning neurovascular coupling, and ability to respond to oxygen demands. Another study has linked antipsychotic related cortical thinning to dopamine D2 receptors by showing that mice treated with antipsychotics have similarly reduced gray matter volume as D2 receptor knock-out mice (14). Exposing the D2 knock-out mice to antipsychotics did not lead to further reductions in gray matter, suggesting that D2 receptors may also play a role in this phenomenon. In addition to the D2 receptor, antipsychotics have affinity to a host of receptors including the serotonin receptors and also affect physiology of the brain (1). However, the role of these features in antipsychotic related cortical thinning has not been thoroughly examined.

Another well documented aspect of antipsychotic related cortical thinning is its regional heterogeneity (10). Overall, antipsychotic medications appear to have more pronounced effects on the prefrontal and temporal cortices compared to other cortical regions and subcortical structures. Here, we make the assumption that these regions are more susceptible because of some underlying molecular, vascular, physiological or functional feature(s). Given the prior evidence and hypotheses suggesting a link with underlying vascular and metabolic capacity and dopamine receptors, it is plausible that regions most vulnerable to antipsychotic induced cortical thinning are enriched/depleted in some of these features. Linking antipsychotic induced cortical thinning with underlying normative features of the brain might thus give clues as to why antipsychotics lead to cortical thinning. This could have further implications for clinical practice and development of novel treatments. However, this link has never been systematically studied in humans.

In this study, we systematically examined what underlying features are associated with the effects of antipsychotic medication on cortical thickness. These analyses were performed across multiple levels of description, from molecular mechanisms, metabolic and physiological features to whole-brain functional patterns in two independent samples. The discovery sample consisted of 131 early psychosis individuals. We measured the effect of lifetime antipsychotic exposure on cortical thickness and then compared the topography of antipsychotic related cortical thinning with normative structural, molecular, physiological and functional features of the cortex. The findings from the discovery sample were subjected to a replication analysis using results from a large meta-analysis (N≥2168) by the Enhancing NeuroImaging Genetics through Meta Analysis (ENIGMA) Consortium (10). Since our findings indicated that antipsychotic related cortical thinning is more pronounced in regions subserving higher-order cognition, we finally considered the associations between antipsychotic related cortical thinning and distribution of cognitive functions across the cortex.

## Results

### Higher lifetime antipsychotic exposure is associated with thinner cortex in the discovery sample

We first sought to replicate previous findings of antipsychotic related cortical thinning in the single site discovery sample (from hereafter referred to as the Turku sample). This sample consisted of 131 individuals either at clinical high risk for psychosis or during first-episode of psychosis. We found that, higher lifetime antipsychotic exposure was associated with thinner mean cerebral cortex when covarying for age, sex and group (clinical high risk or first-episode psychosis) (regression coefficient = −0.00000113, t-value = −4.159, p-value = 0.000059). That is, exposure to about 90 g of chlorpromazine equivalent antipsychotic medication was associated with 0.1 mm thinner cortex. In the sensitivity analyses shown in **supplementary table 1**, lifetime antipsychotic exposure remained statistically significantly associated with mean cortical thickness while covarying for indicators of illness severity. **Figure 1** shows vertex-wise analysis of the association between lifetime antipsychotic exposure and cortical thickness while controlling for age, sex, and group. Cortical thinning was most prominent in the prefrontal, parietal and cingulate cortices. No region showed statistically significant increase in cortical thickness. **Supplementary** Figure 1 shows the vertex-wise results using a more stringent cluster-forming threshold of p = 0.01.

**Figure 1.**
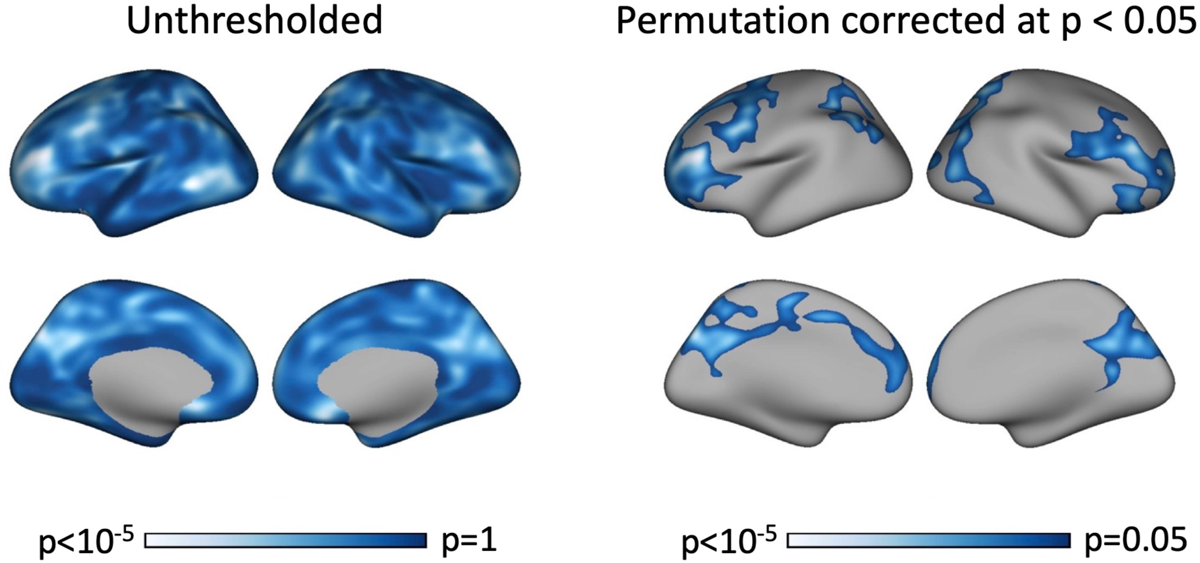
Higher lifetime antipsychotic exposure is associated with lower cortical thickness in the Turku sample (N=131). Figure on the left shows unthresholded results. Figure on the right shows permutation corrected map at p < 0.05. Model includes lifetime antipsychotic exposure, age, sex, and diagnostic group as predictors. Colorbar indicates p-value. There were no vertices where the thickness increased statistically significantly.

### Sensitivity to antipsychotic exposure is associated with molecular, physiological and functional features of the brain in the discovery sample

To understand what underlying biological features are associated with the observed antipsychotic related cortical thinning in the Turku sample, we systematically compared the topographies of the cortical thinning to normative features of the brain (for a full description of all normative brain phenotypes, see Methods). Figure 2 shows the associations between antipsychotic related cortical thinning and the underlying neurobiological features in the Turku sample. Exact values shown in **supplementary table 2**. We discovered 18 features that were associated with antipsychotic related cortical thinning. All associations survived false discovery rate correction for multiple comparisons. Positively correlated features were serotonin 5HT-2A, and 5HT-4 receptors, nicotinic α4β2* receptors, cannabinoid receptor 1, μ-opioid receptors, fMRI functional gradient, neurophysiological measures of delta, low and high gamma, theta power, and intrinsic time scale, synaptic vesicles and cerebral metabolic rate for glucose. That is, regions that typically exhibit high levels of these features (e.g. high serotonin 5HT-2A receptor levels) also had more antipsychotic related cortical thinning. Negatively associated features were serotonin transporter (5-HTT), vesicular acetylcholine transporter, alpha power, T1/T2 ratio and cerebral blood volume. Collectively, analysis in the discovery sample suggested that multiple scales of biological mechanisms (serotonin and cholinergic systems, metabolism, functional networks, neural oscillatory dynamics, synaptic density, and myelination) may underlie the effect of antipsychotics on brain morphology. Several features were measured for more than one tracer. Correlation between antipsychotic related cortical thinning and these alternative tracers are shown in **supplementary** figure 2 and **supplementary table 3**.

**Figure 2.**
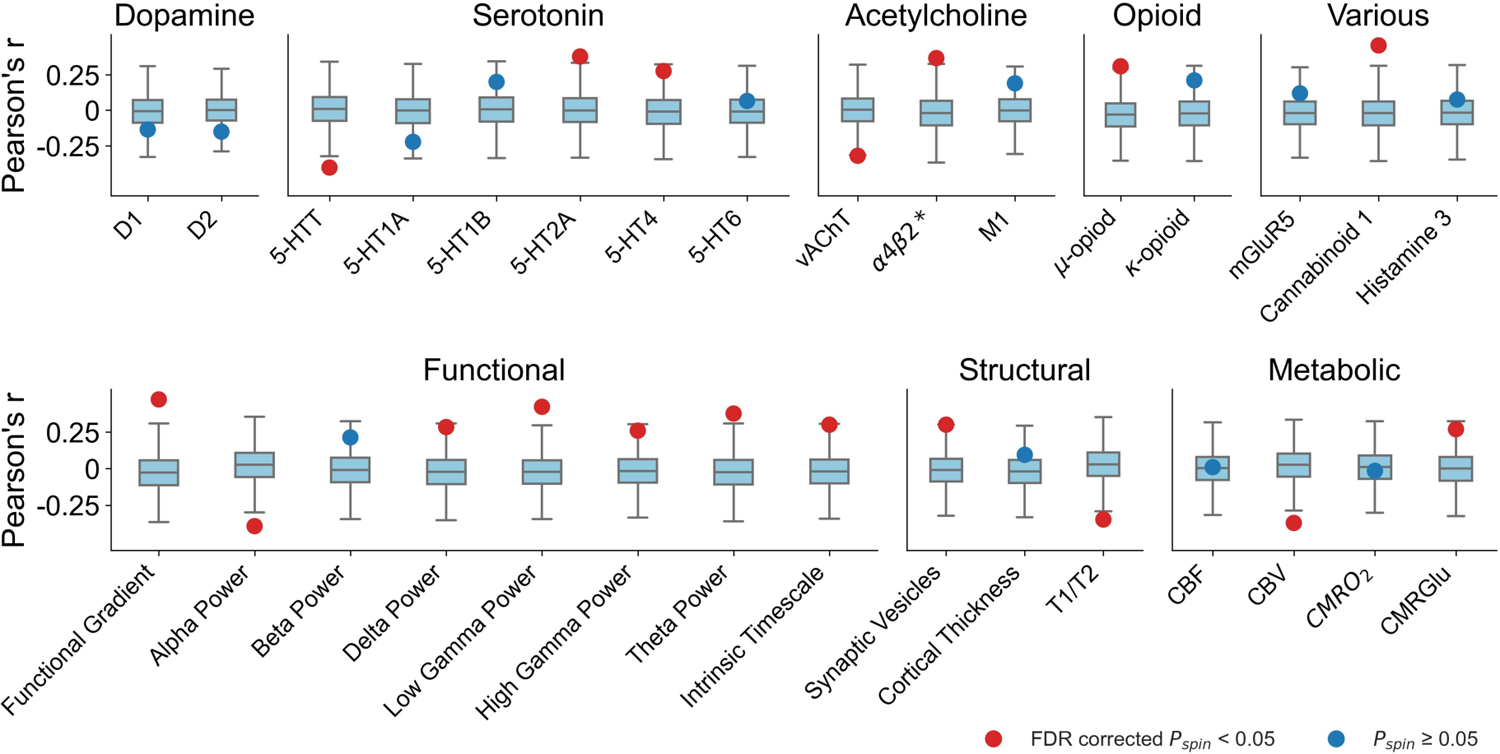
Regional sensitivity to antipsychotic exposure and underlying brain features in the Turku sample. Features associated with antipsychotic related cortical thinning include serotonergic, cholinergic, structural, functional, and metabolic features. Positive correlation indicates that regions that have a higher value of the measured brain feature are more susceptible to the effects of antipsychotics than regions that have a lower value and vice versa. The dots show Pearson’s correlation coefficient r between antipsychotics effects and a given brain feature, the color of the dot indicates statistical significance, all associations survived false discovery rate (FDR) correction for multiple comparisons The ends of the boxes represent the first and third quartiles and the center line represents the median of the null distribution (10,000 rotations), the whiskers represent the non-outlier end-points of the distribution

### Antipsychotic related cortical thinning and molecular, functional, and physiological features in the ENIGMA sample

To replicate the findings of the Turku sample, we used a meta-analysis conducted as a part of a study by the ENIGMA Consortium (hereafter referred to as ENIGMA sample) (10). Shown in Figure 3, we first compared antipsychotic related cortical thinning in the discovery and replication samples to assess comparability. Since antipsychotic related cortical thinning was similar in the two samples (Pearson’s r = 0.47, pspin = 0.0008), we then tested whether our findings from the discovery sample would replicate in the ENIGMA sample. Figure 4 shows that of the 18 associations found in the discovery sample, 14 were replicated. Exact values shown in **supplementary table 4**. 5-HTT, 5-HT2A, 5-HT4, nicotinic α4β2* receptors, cannabinoid 1 receptors, μ-opioid receptors, fMRI functional gradient and interindividual variability, theta, low gamma, and alpha power, T1/T2 ratio, synaptic density and cerebral blood volume. All associations survived false discovery rate correction for multiple comparisons. The replication analysis confirmed that antipsychotic related cortical thinning associates with multiple biological mechanisms, including the serotonin system, and functional networks and neural oscillatory power distributions typical for regions involved in higher cognitive functions. Associations between replicated features were explored in **supplementary** figure 3.

**Figure 3.**
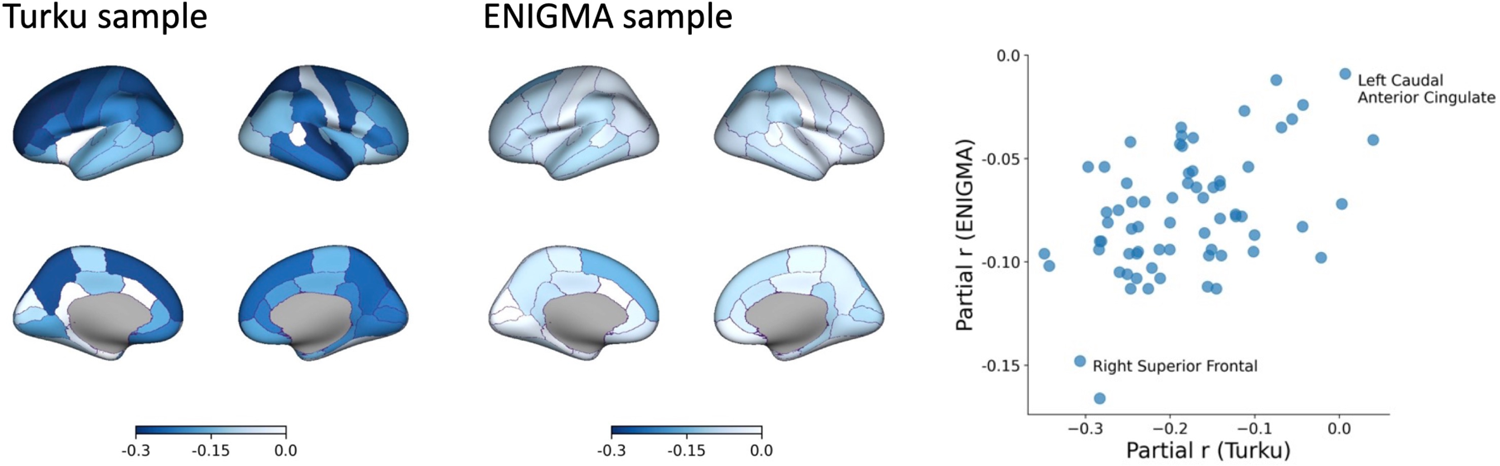
Parcel-wise associations between antipsychotic medication and cortical thickness in the Turku and ENIGMA samples. In the Turku sample, mean cortical thickness in each parcel and lifetime antipsychotic exposure are correlated while controlling for age, sex, and group. In the ENIGMA sample, correlation between mean cortical thickness in each parcel and current antipsychotic dose is calculated while controlling for age and sex. Colorbar indicates partial correlation r. Spatial correlation of antipsychotic effects is similar between the two samples (Pearson’s r=0.47, pspin = 0.0008). Two regions with high and low sensitivity to antipsychotics are labeled in the scatter plot.

**Figure 4.**
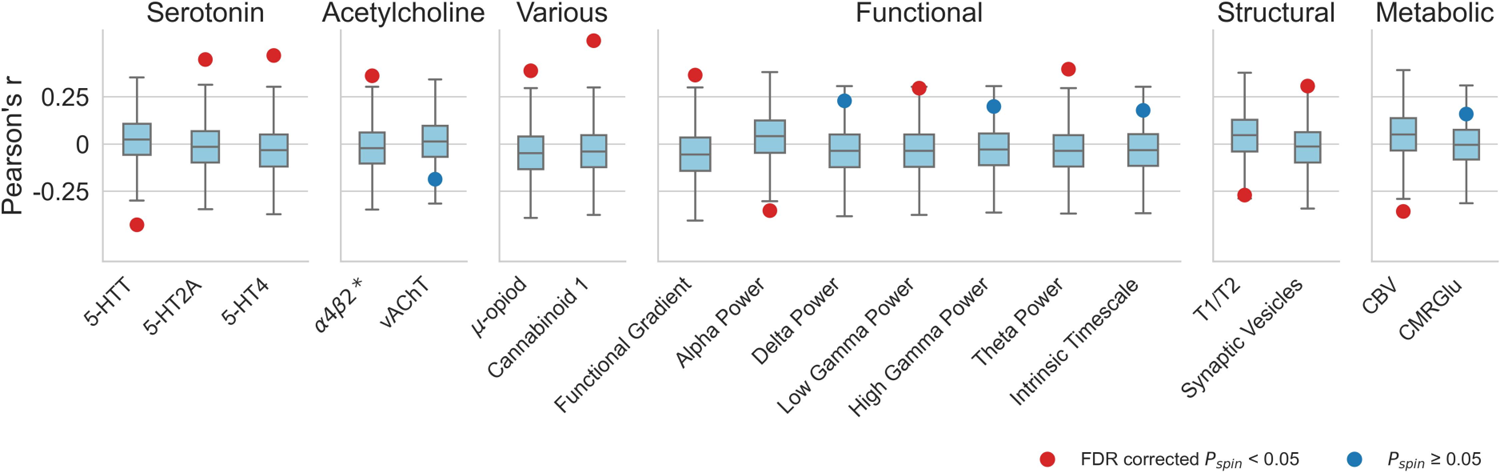
Associations between antipsychotic related cortical thinning and molecular, functional, structural, and metabolic features replicated in the ENIGMA sample. The dots show Pearson’s correlation coefficient r between antipsychotics effects and a given brain feature, the color of the dot indicates statistical significance, all associations survived false discovery rate (FDR) correction for multiple comparisons The ends of the boxes represent the first and third quartiles and the center line represents the median of the null distribution (10,000 rotations), the whiskers represent the non-outlier end-points of the distribution

### Antipsychotic related cortical thinning occurs in regions involved in higher cognitive functions

Finally, we considered potential functional associations, which might link antipsychotic related cortical thinning to the behavioral effects of antipsychotic use. Our analyses thus far have indicated that antipsychotic related cortical thinning may be more pronounced in regions involved in higher cognitive functions. Therefore, we expected antipsychotic related cortical thinning to correlate positively with fMRI activation of higher cognitive functions and negatively with activations during somatosensory tasks. In Figure 5, we show associations between antipsychotic related cortical thinning and different fMRI-derived brain activation patterns, organized into cognitive categories. Altogether, brain regions involved in motivation and executive/cognitive control demonstrate reproducible correlations with antipsychotic related cortical thinning, whereas regions involved in perception are more robust to antipsychotics.

**Figure 5.**
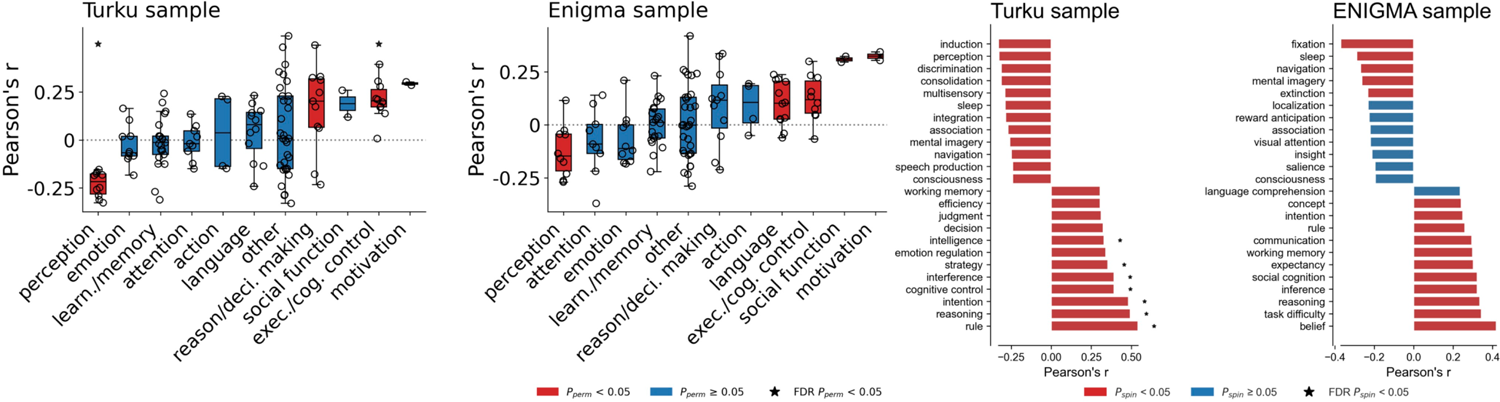
Antipsychotic related cortical thinning and cognitive functions. Probabilistic maps for 123 different cognitive functions were derived from Neurosynth database and correlated with antipsychotic related cortical thinning. In the Figure 5A, the cognitive functions were grouped into 11 categories and ranked based on average correlation. In the Turku sample, we found that functions related to ‘perception’ were negatively correlated and ‘reasoning / decision making’, ‘executive / cognitive control’, and ‘motivation’ were positively correlated with antipsychotic related cortical thinning (two-sided permutation test over groupings). Similarly, in the ENIGMA sample, ‘perception’, ‘executive / cognitive control’, and ‘motivation’, but also ‘language’ and ‘social function’ were associated with antipsychotic related cortical thinning. Top and bottom 10 % of the strongest individual term correlations in both samples are shown in the Figure 5B.

## Discussion

Here, we examined underlying biological features that may underlie sensitivity to antipsychotic related cortical thinning. Using independent discovery and replication samples, we systematically compared topographies of antipsychotic related cortical thinning to a suite of normative features of the brain. Overall, our results show that multiple scales of biological mechanisms from molecular processes to large scale functional gradients are involved in this phenomenon. Notably, we discovered that the serotonin system may play a previously unappreciated role in antipsychotic related cortical thinning. Additionally, acetylcholine, cannabinoid, and opioid neurotransmitter systems are possibly involved. We also found that large scale functional networks and neural oscillatory power distributions render certain brain regions more sensitive to antipsychotics. Finally, in exploratory analyses linking probabilistic cognitive maps from Neurosynth meta-analyses to antipsychotic related cortical thinning, we found that cognitive functions belonging to categories of motivation, and executive and cognitive control were positively associated with antipsychotic related cortical thinning whereas cognitive functions grouped into perception category had a negative association with cortical thinning. Our finding that the serotonin system may underlie antipsychotic related cortical thinning is in line with the essential role that serotonin plays in cortical plasticity and remodeling (15).

Serotonin increases neurite extension, dendritic stabilization and generation of new synapses, whereas depletion of serotonin leads to loss of synapses (16). Serotonin also plays a direct role in cortical thickness highlighted by studies showing that selective serotonin reuptake inhibitors increase cortical thickness (17). Furthermore, serotonin transporter density is the strongest contributor to cortical thinning seen in bipolar, obsessive-compulsive disorder, and schizophrenia (18). Since second generation antipsychotics are potent 5-HT2A antagonists, it is of particular interest that, psilocybin, a 5-HT2A agonist, leads to increase in synaptic density (19). Second generation antipsychotics have also a varying degree of affinity for serotonin receptors making it plausible that part of the cortical remodeling is mediated via the serotonin system. Yet, first generation antipsychotics do not generally bind to serotonin receptors (or have a lower affinity), but they have similar or potentially even larger effect on cortical thickness than second generation antipsychotics (9). Thus, there are probably other mechanisms contributing to the antipsychotic related cortical thinning.

Accordingly, we also found that the principal functional gradient, low gamma and theta power are positively associated antipsychotic related cortical thinning. The principal functional gradient delineates primary unimodal sensorimotor regions from higher-order transmodal regions involved in cognitive functions (20). Our finding thus indicates that cortical thickness in regions involved in higher cognition is particularly sensitive to antipsychotics. On the other hand, regions involved in unimodal sensory or motor functions is relatively spared. One reason for the sensitivity of higher-order transmodal brain regions for antipsychotics might be that these regions continue to develop longer than sensorimotor regions, retaining their plasticity late into adolescence and early adulthood (21). Antipsychotics are often first prescribed when these sensitivity windows still remain open, which could explain why there is more pronounced remodeling. On the other hand, unimodal regions are under a tighter genetic influence and may not be as amenable to cortical modeling (22). Interestingly, the transmodal regions also show the highest energy signaling cost (23), which might make these regions particularly vulnerable for disturbances in cerebral blood flow. Previous studies have shown that antipsychotics induce changes in cerebral blood flow (13) and we found that regions with lower cerebral blood volume is associated with increased antipsychotic related cortical thinning. Altogether these findings lend some support to the hypothesis that antipsychotics may push the cortex into an unsustainable state that leads to cortical remodeling specifically in regions with high energy demand and low blood supply (13). Correlations with MEG power bands complement the finding with resting state measures. Theta and gamma power, that show positive correlation with antipsychotic related cortical thinning, are associated with higher cognitive functions (24,25). On the other hand, a negative association was found with the alpha band power that is almost exclusively limited to the posterior and visual regions (26). Thus, the overall emerging picture from this study is that antipsychotics lead to remodeling of those cortical regions that are involved in higher cognitive functions.

Our analyses associating probabilistic cognitive maps from Neurosynth meta-analyses with antipsychotic related cortical thinning further suggest that antipsychotics target regions involved in higher cognition. Research thus far has not associated antipsychotic related cortical thinning unequivocally with neither poorer nor better outcomes (11). Generally antipsychotics, and particularly second generation, may have weakly beneficial effects on cognitive deficits of schizophrenia (27). However, the effect on executive function may be more variable (28). Some reports (29), but not all (30) have also suggested that antipsychotic use is associated with lower motivation. We speculate that some of the beneficial as well as harmful consequences of antipsychotic use may be mediated via cortical remodeling.

Higher levels of nicotinic α4β2* receptors, which antipsychotics inhibit, were associated with cortical thinning in both discovery and replication sample. Acetylcholine is involved in cortical maturation and plasticity and thus some effects of antipsychotics are potentially mediated through this neurotransmitter system as well (31). We also found that cannabinoid 1 receptors, and μ-opioid receptors were associated with this phenomenon. However, since antipsychotics do not directly bind to cannabinoid 1 or μ-opioid receptors the role of these receptors remains to be clarified. All currently used antipsychotics act on dopamine D2 receptors. Yet, dopamine D2 or D1 receptor levels were not associated with antipsychotic related cortical thinning.

In this study, we used a discovery sample to narrow down the search space and then a replication sample to confirm our findings. While the antipsychotic related cortical thinning in discovery and replication samples had a moderately positive correlation, there are a number of notable differences in these samples. The discovery sample is a relatively small, single site study with a homogenous catchment area using a single scanner. The participants individuals at early psychosis and mostly treated with second-generation antipsychotic. Cortical thickness was related to total lifetime antipsychotic exposure, determined retrospectively for each participant from electronic health records. Importantly, sensitivity analyses in this sample indicated that cortical thinning is better explained by antipsychotic exposure than severity of illness. On the other hand, in the large sample ENIGMA study, patients were diagnosed with schizophrenia, were scanned at multiple locations, with various scanners and protocols and cortical thickness was correlated with current dose rather than exposure as in the discovery sample. Yet, despite many differences between the discovery and replication samples, most features were still replicated. On the whole, these results show that multiple scales of biological mechanisms underlie the effect of antipsychotics on brain morphology.

## Methods

### Discovery sample collected in Turku, Finland Participants

All participants were recruited from psychiatric services of the Hospital District of Southwest Finland and scanned at Turku PET Centre using Philips Ingenuity 3T MRI scanner (Philips Healthcare, The Netherlands). Details of the recruitment are described elsewhere (32,33). Participants with an IQ under 70, a significant somatic or neurological illness that might affect brain structure or function, earlier head injury with loss of consciousness for over five minutes, or alcohol dependence during the preceding 6 months were also excluded. All images were read by a neuroradiologist and individuals with significant findings were excluded. The intent-to-study group consisted of age- and sex-matched 87 first-episode psychosis patients (FEP), 56 clinical high-risk for psychosis patients (CHR) between 18-50 years of age. Written informed assent and consent were obtained from all the participants. The study protocol was approved by the Ethics Committee of the Hospital District of Southwest Finland and the study was conducted in accordance with the Declaration of Helsinki.

From the intent-to-study group, one CHR patient was excluded for having incomplete records of past antipsychotic use. Five FEP’s and six more CHR’s were excluded for either motion during the scan, radiological findings, or technical issues during acquisition resulting in unusable T1.

**Table 1** shows the demographic and clinical characteristics of the 131 individuals included in the final discovery sample. Four first-episode psychosis patients (4.9 %) and 22 clinical high-risk individuals (44.9 %) were antipsychotic naive, and the rest of the sample were exposed to antipsychotics. Average daily dose at the time of scanning in those who were taking antipsychotics was 321.84 (± 233.09) mg chlorpromazine in the first-episode psychosis group and 167.06 (±129.33) mg chlorpromazine in the high-risk group. **Supplementary table 5** lists all antipsychotic medications ever used in this sample and the number of people who were on them at any point of their lives.

**Table 1.** shows the demographics of the discovery sample collected at Turku, Finland.

|  | CHR (N = 49) | FEP (N = 82) |
| --- | --- | --- |
| Age | 25.75 (6.09) | 26.76 (5.98) |
| Males (%) | 28 (57.1) | 47 (57.3) |
| Lifetime antipsychotic exposure | 7733.04 (18902.10) | 21199.03 (25207.65) |
| Current antipsychotic dose | 57.96 (109.69) | 290.44 (241.24) |
| Total symptom score | 35.55 (10.09) | 38.04 (12.29) |
| Positive symptoms | 13.11 (4.04) | 16.85 (6.33) |
| Negative symptoms | 9.19 (3.37) | 9.41 (3.73) |
| GAF | 53.80 (11.38) | 51.38 (14.99) |
| SOFAS | 53.37 (11.66) | 48.59 (15.12) |
| Days in Hospital | 18.56 (30.52) | 44.24 (48.15) |
| Times admitted to Hospital | 0.62 (0.70) | 1.07 (0.75) |
| DOI | - | 1.37 (2.59) |
| Antipsychotic naive (%) |  |  |
| No | 27 (55.1) | 78 (95.1) |
| Yes | 22 (44.9) | 4 (4.9) |
| Currently antipsychotic free (%) |  |  |
| No | 17 (34.7) | 74 (90.2) |
| Yes | 32 (65.3) | 8 (9.8) |

CHR=Clinical High Risk for psychosis as determined by SIPS/SOPS interview, FEP=First Episode Psychosis, GAF=Global Assessment of Functioning, SOFAS=Social and Occupational Functioning Assessment Scale, DOI=Duration of Illness. Table presents means and standard deviations unless otherwise stated.

### Lifetime antipsychotic exposure

Lifetime antipsychotic exposure was calculated from electronic medical records of the psychiatric services of the Hospital District of Southwest Finland. **Supplementary table 5** shows all antipsychotics that were used in this sample. Most commonly used antipsychotics were risperidone, quetiapine, olanzapine and aripiprazole. None of the patients were and clozapine. All medications ever prescribed to the participants were recorded, converted to chlorpromazine equivalents and added together. Medications given on an as-needed basis were not included.

### Clinical assessments

The measures used are described in detail elsewhere. Briefly, diagnoses were confirmed using the Structured Clinical Interview for DSM-IV disorders (SCID-I/NP) and clinical high-risk status using the SIPS/SOPS (34). assessed using either the Positive and Negative Symptom Scale (SCI-PANSS) or the Brief Psychiatric Rating Scale (BPRS) (24-items, version 4.0)(35,36). The PANSS scores were converted to correspond the BPRS 18-item scores and the BPRS 24-item scores reduced to correspond the BPRS 18-item scores. Thus, in the following analyses, the total symptom score refers to the sum of those 18 items, positive symptoms score to the sum of the 8 positive symptom items and negative symptoms score to the sum of the 5 negative symptom items. All items were rated on a scale of 1 to 7. Global Assessment of Functioning (GAF) scale, and Social and Occupational Functioning Assessment Scale (SOFAS) were used to assess functioning (37,38). Along with prescribed antipsychotic medications, we also recorded the number of days each participant had been an inpatient in a mental health hospital and the number of times they were admitted to a mental health hospital as an inpatient. Symptom scores, measures of function, number of days in a mental health hospital, and number of times admitted to a mental health hospital were used as proxies of illness severity.

### MRI acquisition and processing

All participants underwent an MRI scan with a Philips Ingenuity TF 3-Tesla PET/MR scanner. T1-weighted (Ultrafast Gradient Echo 3D, TR = 8.1 ms, TE-time = 3.7 ms, flip angle 7°, FOV = 256×256×176 mm^3^ and voxel size 1×1×1 mm^3^) image was collected from all subjects. T1 images were preprocessed using the longitudinal pipeline in Freesurfer v7.1.1 (39). All scans and reconstructed images were visually inspected and reconstructed surfaces manually corrected as needed by R-LA.

### Statistical analyses

We first estimated the effect of lifetime antipsychotic exposure on total cortical thickness. The average cortical thickness across the whole cortical mantle was used as the outcome measure and lifetime antipsychotic exposure, age, sex, and group as explanatory variables in a linear regression model: mean cortical thickness ∼ intercept + β1 (lifetime antipsychotic exposure) + β2 (age) + β3 (sex) + β4 (group)+ ε. Robust MM-estimation method in robustbase library running on R (v4.2.2) was used due to its insensitivity to outliers (40). Since, it is possible that illness severity could explain both higher antipsychotic exposure and lower cortical thickness, we conducted a series of sensitivity analyses using indices of overall symptom or illness severity as covariates in the regression model. These sensitivity analyses were carried out by including one of the following measures at a time as an additional covariate in the above-described regression analyses: total symptom score, GAF, SOFAS, total number of days spent in a mental health hospital, or the number of times admitted to a mental health hospital.

Regional effects of lifetime antipsychotic exposure on cortical thickness were estimated using vertex-wise analyses, carried out using glm-fit in Freesurfer 7.1.1. In this analysis, vertex-wise cortical thickness was predicted using lifetime antipsychotic exposure with age, sex and group as covariates. We used 1000 permutations to correct for multiple comparisons with cluster-wise p-value of 0.05 and cluster forming p-values of 0.05, and 0.01 in a more stringent analysis. Bonferroni correction was used to take into account that tests were conducted in two hemispheres.

### An independent replication data set from an ENIGMA study

To replicate the findings from the discovery sample, we used data from an ENIGMA (Enhancing Neuroimaging Genetics through Meta-Analysis) consortium study on cortical thickness in schizophrenia (10). In that study, for a subset of patients, antipsychotic medication dose at the time of the scanning was available. Of those, 2236 (66%) were on second-generation (atypical) antipsychotic, 447 (13%) on first-generation (typical), and 265 (8%) on both second-generation and first-generation antipsychotic. Current antipsychotic dose was converted to chlorpromazine equivalents based on Woods’ calculations (www.scottwilliamwoods.com/files/Equivtext.doc). After quality control, partial correlations between current chlorpromazine equivalent dose and cortical thickness were calculated for each Desikan-Killiany parcel while controlling for age and sex in 2168-2175 individuals depending on the parcel.

### Testing the association between underlying cortical structural and functional features and antipsychotic related cortical thinning

Associations between regional antipsychotic effects on cortical thickness and regional differences in underlying features of the brain were tested using the neuromaps toolbox (41). For the discovery analyses, we selected a total of 33 unique normative structural and functional features of cortical organization provided as a part of the neuromaps toolbox (**Supplementary table 6**). These included neuroreceptor maps for dopamine, serotonin, acetylcholine, glutamate, GABA, cannabinoid, histamine, and opioid systems, measures of blood flow, metabolism, average cortical thickness, myelination, synaptic density, resting state functional MRI connectivity measures of interindividual variability and unimodal-transmodal functional gradient, and MEG-derived neural oscillatory power distributions for six canonical frequency bands (alpha-high gamma) from published studies (18). Six receptors/transporters were measured with more than one positron emission tomography (PET) tracer. These duplicate neuroreceptor maps were included in a supplementary analysis for the sake of completeness. We omitted PET measures of cortical dopamine transporter as it is not clear if these tracers are sensitive enough in the cortex.

The vertex-wise map of the effects of lifetime antipsychotic exposure on cortical thickness from the Turku sample was first parcellated using the Desikan-Killiany cortical atlas (42). We chose this parcellation to be consistent with the ENIGMA data which is only provided with this parcellation. Selected feature annotations provided by the neuromaps were also parcellated using the same atlas. Pairwise Pearson’s correlation r between the parcellated lifetime antipsychotic exposure and parcellated brain features were calculated and statistical significance was assessed using a conservative null model that preserves the spatial-autocorrelation of brain maps (“spin test”)(43,44). In the spin test, one of the pairs in each pairwise comparison, parcel coordinates are projected onto a spherical surface and then randomly rotated. Original parcels are reassigned the value of the closest rotated parcel and the process is repeated 10,000 times. Correlations between the non-rotated pair of the comparison and each permuted map are calculated and the original Pearson’s correlation r is compared against this null distribution allowing the calculation of permutation p-value (pspin). In the spin test, a null distribution of correlations is constructed by randomly rotating one of the brain maps used in a pairwise correlation and repeating the correlation (10,000 repetitions). The empirical correlation is compared against this null distribution to calculate a permutation p-value (pspin).

For the replication analyses using the ENIGMA sample, we selected only the features that were statistically significantly (pspin < 0.05) associated with the effects of lifetime antipsychotic exposure in the discovery sample. Associations between the ENIGMA sample and the features were tested as above and then corrected for multiple comparisons using false discovery rate correction.

### Cognitive functions and antipsychotic related cortical thinning

As a follow-up analysis and in order to further interrogate the functional relevance of our primary analyses, we explored potential links between antipsychotic related cortical thinning and various cognitive functions with the assumption that antipsychotic-related cortical thinning would be more prominent in regions that are involved in higher cognitive functions. To this end, we used the Neurosynth database (neurosynth.org), which includes probabilistic brain activation/deactivation maps from over 14,000 fMRI studies, to conduct voxel-wise meta-analyses for 123 cognitive terms from the Cognitive Atlas (45,46). These terms include umbrella terms (eg. “attention”, “emotion”), specific cognitive processes (eg. “visual attention”, “episodic memory”), behaviors (eg. “eating”, “sleep”), and emotional states (eg. “fear”, “anxiety”). The terms were then grouped into 11 categories (“Action”, “Learning and Memory”, “Emotion”, “Attention”, “Reasoning and Decision Making”, “Executive/Cognitive control”, “Social Function”, “Perception”, “Motivation”, “Language”, and “other”) (http://www.cognitiveatlas.org/concepts/categories/all)(46). The probabilistic meta-analytical maps were parcellated using the Desikan-Killiany atlas and their association with antipsychotic related cortical thinning was tested in both the discovery sample and replication data using spin tests as above. To test whether the categories on average were associated with antipsychotic related cortical thinning we conducted a two-sided permutation test where the average category correlation was compared against a null distribution obtained by permuting the category membership 10,000 times. In order to gain further insights into antipsychotic related cortical thinning and cognitive functions, we also explored the top and bottom 10% of the individual term correlations in both datasets.

## Acknowledgements

Collection of the Turku sample was supported by grants to Raimo K.R. Salokangas, Jarmo Hietala and Nikolaos Koutsouleris.

## Code and data availability

Group level statistical maps, parcel-wise partial correlation values and code for validation and replication of all neuromaps analyses and figures is available online: https://github.com/ltuominen/AP_brainorganization. Individual-level data cannot be shared.

