## supplemental material for "Molecular, physiological and functional features underlying cortical thinning related to antipsychotic medication use"

Supplementary figures and tables for Tuominen et al., Molecular, physiological and functional features underlying cortical thinning related to antipsychotic medication use

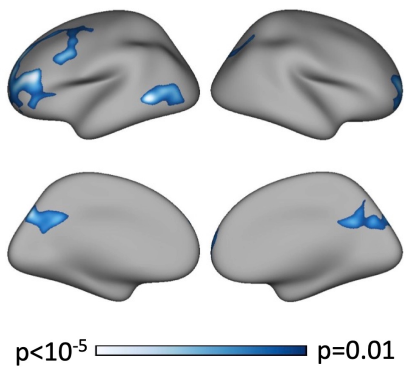


**Supplementary Figure 1 Association between lifetime antipsychotic exposure and cortical thickness thresholded at p=0.01.** Model includes age, sex, and diagnostic group as nuisance variables. Colorbar indicates p-value. Results are permutation corrected for multiple comparisons.

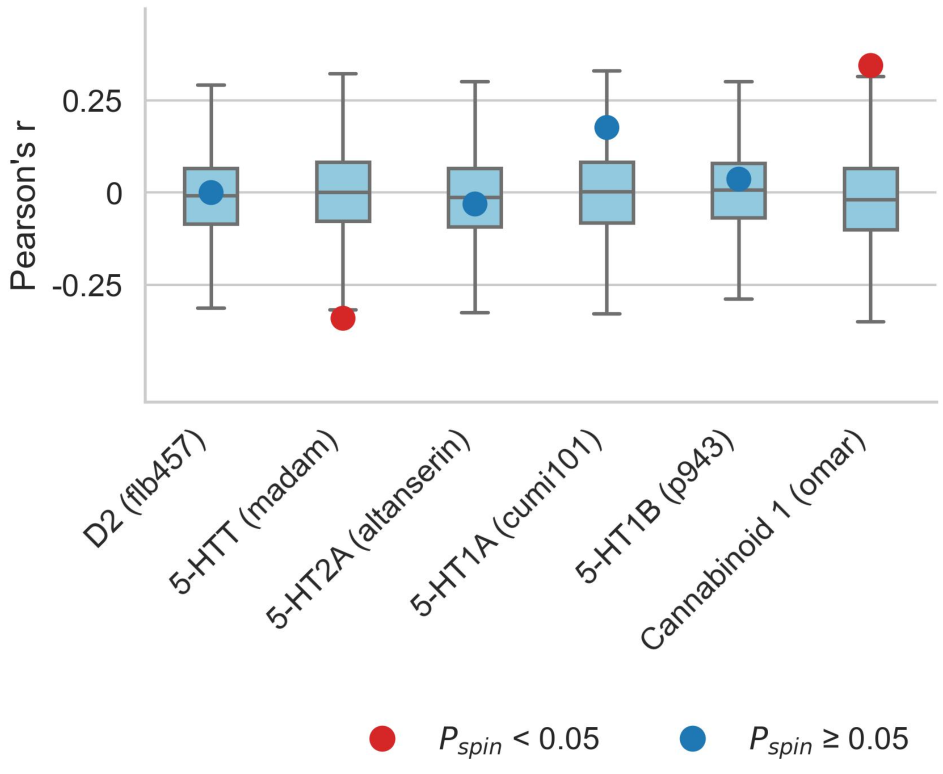


**Supplementary Figure 2 Antipsychotic sensitivity and brain organization with additional measures.** Six features of brain organization were measured with more than one tracer. For the sake of completeness, these measures were also correlated with AP effects on cortical thickness in the Turku sample. In line with the analysis using the primary measures for these features, no statistically significant associations were found.

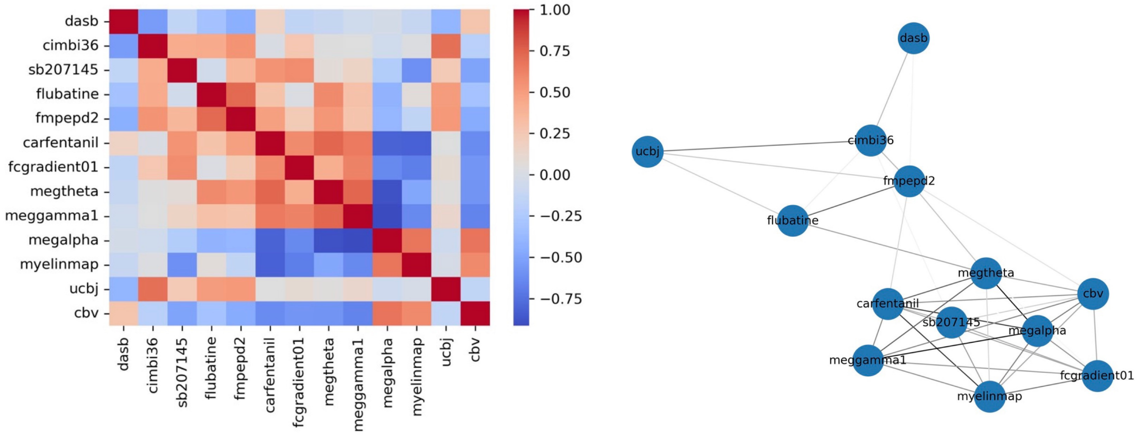


**Supplementary Figure 3** Correlations between features of the brain that are associated with antipsychotic related cortical thinning. Left panel show correlation matrix, colorbar indicates Pearson correlation coefficient r. Right panel shows a spring-embedded representation of the correlations. Topographies of serotonin transporter, synaptic density, 5-HT2A, CB1, and $\alpha4\beta2$* receptors appear clustered, another cluster is formed by functional measures, cortical myelin, cerebral blood flow, $\mu$- opioid and 5-HT4 receptors.

**Supplementary table 1 Sensitivity analyses**. The table shows the estimate, standard error, t-value and p-value of antipsychotic exposure on average cortical thickness when including one of the covariates at a time in the model.

| Covariate | Estimate | Std. Error | t-value | p-value |
|---|---|---|---|---|
| **Total symptoms score** | -1.2E-06 | 2.5E-07 | -4.77 | 0.0000053 |
| **Positive symptoms score** | -1.2E-06 | 2.6E-07 | -4.60 | 0.0000106 |
| **Negative symptoms score** | -1.2E-06 | 2.5E-07 | -4.86 | 0.0000036 |
| **BMI** | -1.1E-06 | 2.5E-07 | -4.30 | 0.0000347 |
| **Hospital days** | -1.0E-06 | 3.1E-07 | -3.31 | 0.0012269 |
| **Times admitted** | -1.1E-06 | 2.8E-07 | -3.72 | 0.0003040 |
| **GAF** | -1.2E-06 | 2.8E-07 | -4.12 | 0.0000688 |
| **SOFAS** | -1.2E-06 | 2.8E-07 | -4.19 | 0.0000532 |

**Supplementary table 2** shows correlations between antipsychotic related cortical thinning and normative features of the brain in the discovery sample (same data as in figure 2)

| tracer / measure | class | target | rho | pspin | fdr_corrected_p_value |
|---|---|---|---|---|---|
| **sch23390** | dopamine | D1 | -0.13611309816019400 | 0.2522747725227480 | 0.3128207179282070 |
| **fallypride** | dopamine | D2 | -0.15145839192235800 | 0.1648835116488350 | 0.21297453587974500 |
| **dasb** | serotonin | 5-HTT | -0.4041914204738630 | 0.0017998200179982 | 0.007970631508277740 |
| **way100635** | serotonin | 5-HT1A | -0.2230886044601000 | 0.06729327067293270 | 0.1097942837295220 |
| **az10419369** | serotonin | 5-HT1B | 0.19896995442264000 | 0.1095890410958900 | 0.1477069684335910 |
| **cimbi36** | serotonin | 5-HT2A | 0.377865933841217 | 0.0011998800119988000 | 0.007439256074392560 |
| **sb207145** | serotonin | 5-HT4 | 0.27514812603014100 | 0.018998100189981000 | 0.0368088191180882 |
| **gsk215083** | serotonin | 5-HT6 | 0.06486864703669620 | 0.593040695930407 | 0.6339400542704350 |
| **feobv** | acetylcholine | vAChT | -0.321011588836917 | 0.006299370062993700 | 0.017752770177527700 |
| **flubatine** | acetylcholine | alpha4 beta2* | 0.3662865373866890 | 0.004899510048995100 | 0.015498450154984500 |
| **lsn3172176** | acetylcholine | M1 | 0.18914965240899800 | 0.0948905109489051 | 0.1394860513948610 |
| **abp688** | various | mGluR5 | 0.11861066017734100 | 0.32836716328367200 | 0.3915146946843780 |
| **fmpepd2** | various | Cannabinoid 1 | 0.45584978842246300 | 0.000999900009999 | 0.007439256074392560 |
| **gsk189254** | various | Histamine 3 | 0.07465882818090720 | 0.5506449355064490 | 0.6096426071678550 |
| **carfentanil** | opioid | mu-opiod | 0.308922119988958 | 0.014698530146985300 | 0.030376962303769600 |
| **ly2795050** | opioid | kappa-opioid | 0.20964293008754100 | 0.098990100989901 | 0.1394860513948610 |
| **fcgradient01** | functional | Functional Gradient | 0.4736608549761900 | 9.99900009999E-05 | 0.0030996900309969000 |
| **megalpha** | functional | Alpha Power | -0.39369935598264800 | 0.0016998300169983000 | 0.007970631508277740 |
| **megbeta** | functional | Beta Power | 0.21379662504879700 | 0.0842915708429157 | 0.13065193480651900 |
| **megdelta** | functional | Delta Power | 0.28439367925684500 | 0.014298570142985700 | 0.030376962303769600 |
| **meggamma1** | functional | Low Gamma Power | 0.4230748599315880 | 0.0005999400059994000 | 0.007439256074392560 |
| **meggamma2** | functional | High Gamma Power | 0.25983323129290300 | 0.0212978702129787 | 0.0388372927413141 |
| **megtheta** | functional | Theta Power | 0.3769190538769740 | 0.004999500049995000 | 0.015498450154984500 |
| **megtimescale** | functional | Intrinsic Timescale | 0.3007630887583380 | 0.008599140085991400 | 0.022214445222144500 |
| **ucbj** | structural | Synaptic Vesicles | 0.30075589867662100 | 0.009499050094990500 | 0.02265158099574660 |
| **cortical thickness** | structural | Cortical Thickness | 0.09457914724924790 | 0.42295770422957700 | 0.48561810485618100 |
| **myelin** | structural | T1/T2 | -0.34840135738894000 | 0.0038996100389961000 | 0.01511098890110990 |
| **cbf** | metabolic | CBF | 0.009795333069901100 | 0.9342065793420660 | 0.9342065793420660 |
| **cbv** | metabolic | CBV | -0.37187655201844700 | 0.0011998800119988000 | 0.007439256074392560 |
| **cmr02** | metabolic | CMRO2 | -0.014991231074433000 | 0.9032096790320970 | 0.9333166683331670 |
| **cmrglc** | metabolic | CMRGlu | 0.26999789426642900 | 0.022897710228977100 | 0.039434945394349500 |

**Supplementary table 3** shows correlations between antipsychotic related cortical thinning and alternative tracers (same data as in supplementary figure 3)

| tracer | class | target | rho | pspin |
|---|---|---|---|---|
| **flb457** | dopamine | D2 | -5.79697230025378E-05 | 0.9996000399960000 |
| **madam** | serotonin | 5-HTT | -0.34088637208110100 | 0.0040995900409959 |
| **cumi101** | serotonin | 5-HT1A | -0.031009435680068000 | 0.7922207779222080 |
| **p943** | serotonin | 5-HT1B | 0.17632035708924800 | 0.138986101389861 |
| **altanserin** | serotonin | 5-HT2A | 0.036385225482637400 | 0.7381261873812620 |
| **omar** | various | Cannabinoid 1 | 0.34450355205267800 | 0.005099490050994900 |

**Supplementary table 4** shows correlations between antipsychotic related cortical thinning and normative features of the brain in the replication sample (same data as in figure 4)

| tracer / measure | class | target | rho | pspin | fdr_corrected_p_value |
|---|---|---|---|---|---|
| **dasb** | serotonin | 5-HTT | -0.42730137785045700 | 0.00039996000399960000 | 0.0035996400359964000 |
| **cimbi36** | serotonin | 5-HT2A | 0.44723884247094900 | 0.00029997000299970000 | 0.0035996400359964000 |
| **sb207145** | serotonin | 5-HT4 | 0.4685705038863450 | 0.0010998900109989000 | 0.004949505049495050 |
| **flubatine** | acetylcholine | alpha4 beta2* | 0.361200659937138 | 0.0028997100289971 | 0.007456397217421110 |
| **feobv** | acetylcholine | vAChT | -0.18615621847142200 | 0.12138786121387900 | 0.13892360763923600 |
| **fmpepd2** | various | Cannabinoid 1 | 0.5468469740436400 | 0.0005999400059994000 | 0.0035996400359964000 |
| **carfentanil** | various | $\mu$-opiod | 0.38768215355838800 | 0.0025997400259974 | 0.007456397217421110 |
| **fcgradient01** | functional | Functional Gradient | 0.3648219948521910 | 0.006199380061993800 | 0.012398760123987600 |
| **megtimescale** | functional | Intrinsic Timescale | 0.17808098151252700 | 0.16498350164983500 | 0.16498350164983500 |
| **megtheta** | functional | Theta Power | 0.3962061791475740 | 0.0016998300169983000 | 0.006119388061193880 |
| **meggamma2** | functional | High Gamma Power | 0.19868373094673000 | 0.12348765123487700 | 0.13892360763923600 |
| **meggamma1** | functional | Low Gamma Power | 0.2956246730624560 | 0.0188981101889811 | 0.028347165283471700 |
| **megdelta** | functional | Delta Power | 0.22859793901970000 | 0.0822917708229177 | 0.10580370534375100 |
| **megalpha** | functional | Alpha Power | -0.35220229889986500 | 0.0053994600539946000 | 0.012148785121487900 |
| **myelin** | structural | T1/T2 | -0.2702431372254100 | 0.033996600339966 | 0.047072215855337500 |
| **ucbj** | structural | Synaptic Vesicles | 0.3068443681943620 | 0.0091990800919908 | 0.015053040150530400 |
| **cbv** | metabolic | CBV | -0.3566093662360660 | 0.007799220077992200 | 0.014038596140386000 |
| **cmruglu** | metabolic | CMRGlu | 0.15876486313861500 | 0.1643835616438360 | 0.16498350164983500 |

**Supplementary table 5** shows all the antipsychotic medications that were used in the Turku sample and a number of patients that were ever exposed to these antipsychotics.

| | **FEP** | **CHR** |
|---|---|---|
| **Second generation antipsychotics** | | |
| **risperidone (n)** | 50 | 12 |
| **quetiapine (n)** | 35 | 13 |
| **olanzapine (n)** | 31 | 7 |
| **aripiprazole (n)** | 12 | 3 |
| **paliperidone (n)** | 5 | 0 |
| **asenapine (n)** | 4 | 0 |
| **ziprasidone (n)** | 1 | 0 |
| **sulpride (n)** | 1 | 0 |
| **First generation antipsychotics** | | |
| **perphenazine (n)** | 6 | 2 |
| **haloperidol (n)** | 3 | 2 |
| **flupentixol (n)** | 2 | 0 |
| **levomepromazine (n)** | 2 | 0 |

**Supplementary table 6** shows measures of normative structural and functional features of the cortex used in this study.
For more details please see:
https://docs.google.com/spreadsheets/d/1oZecOsvtQEh5pQkIf8cB6CyhPKVrQuko/edit#gid=1162991686

| | Author/Dataset | Year | Tracer / Measure | Target | Reference |
|---|---|---|---|---|---|
| 0 | Jaworska | 2020 | fallypride | D2 | Jaworska et al., 2020, Neuropsychopharm |
| 1 | Kaller | 2017 | sch23390 | D1 | Kaller et al., 2017, Eur J Nucl Med Mol Imaging |
| 2 | Radnakrishnan | 2018 | gsk215083 | 5-HT6 | Radhakrishnan et al., 2018, J Nucl Med |
| 3 | Beliveau | 2017 | cimbi36 | 5-HT2A | Beliveau et al., 2017, J Neurosci |
| 4 | Savli | 2012 | way100635 | 5-HT1A | Savli et al., 2012, Neuroimage |
| 5 | Beliveau | 2017 | az10419369 | 5-HT1B | Beliveau et al., 2017, J Neurosci |
| 6 | Beliveau | 2017 | sb207145 | 5-HT4 | Beliveau et al., 2017, J Neurosci |
| 7 | Fazio | 2016 | madam | 5-HTT | Fazio et al., 2016, Neuroimage |
| 8 | Tuominen | NA | feobv | vAChT | NA |
| 9 | Hillmer | 2016 | flubatine | alpha4 beta2* | Hillmer et al., 2016, Neuroimage |
| 10 | Naganawa | 2020 | lsn3172176 | M1 | Naganawa et al., 2020, J Nucl Med |
| 11 | Margulies | 2016 | fcgradient01 | Functional Gradient | Margulies et al., 2016, PNAS |
| 13 | Hcps | 2022 | megalpha | Alpha Power | Shafiei et al., 2022, Plos Biology |
| 14 | Hcps | 2022 | megdelta | Delta Power | Shafiei et al., 2022, Plos Biology |
| 15 | Hcps | 2022 | megbeta | Beta Power | Shafiei et al., 2022, Plos Biology |
| 16 | Hcps | 2022 | meggamma1 | Low Gamma Power | Shafiei et al., 2022, Plos Biology |
| 17 | Hcps | 2022 | meggamma2 | High Gamma Power | Shafiei et al., 2022, Plos Biology |
| 18 | Hcps | 2022 | megtheta | Theta Power | Shafiei et al., 2022, Plos Biology |
| 19 | Hcps | 2022 | megtimescale | Intrinsic Timescale | Shafiei et al., 2022, Plos Biology |
| 20 | Finnema | 2016 | ucbj | Synaptic Vesicles | Finnema et al., 2018, J Cereb Blood Flow Metab |
| 21 | Hcps | 2016 | thickness | Cortical Thickness | Glasser et al., 2016, Nature |
| 22 | Hcps | 2016 | myelin | T1/T2 | Glasser et al., 2016, Nature |
| 23 | Dukart | 2018 | flumazenil | $GABA_A$ | Dukart et al., 2018, Sci Rep |
| 24 | Dubois | 2015 | abp688 | mGluR5 | Dubois et al., 2016, Eur J Nucl Med Mol Imaging |
| 25 | Laurikainen | 2018 | fmpepd2 | Cannabinoid 1 | Laurikainen et al., 2019, Neuroimage |
| 26 | Vijay | 2018 | ly2795050 | kappa-opioid | Vijay et al., 2018, Neuropsychopharmacology |
| 27 | Kantonen | 2020 | carfentanil | mu-opiod | Kantonen et al., 2020, Neuroimage |
| 28 | Gallezot | 2017 | gsk189254 | Histamine 3 | Gallezot et al., 2010, J Cereb Blood Flow Metab |
| 29 | Raichle | NA | cbf | CBF | Vaishnavi et al., 2010, PNAS |
| 30 | Raichle | NA | cbv | CBV | Vaishnavi et al., 2010, PNAS |
| 31 | Raichle | NA | cmr02 | $CMRO_2$ | Vaishnavi et al., 2010, PNAS |
| 32 | Raichle | NA | cmruglu | CMRGlu | Vaishnavi et al., 2010, PNAS |
